## Supplementary figures and table for "Spatiotemporal model of cellular mechanotransduction via Rho and YAP"

### A Supporting Information

The data used for generating the figures, as well as animations of the concentration profiles, raw concentration profile data, additional plots and MATLAB scripts that can be used to reproduce the data and figures are available online. Here we present some additional results and figures that complement the data in the main text.

#### A.1 Additional information about the model

The harmonic potential we use is defined as

$$\phi(x) = \begin{cases} \frac{\left(x - \frac{x_{\text{nucl front}} + x_{\text{nucl back}}}{2}\right)^2}{x_{\text{nucl back}} - x_{\text{nucl front}}} + \frac{1}{4}\left(x_{\text{nucl front}} - x_{\text{nucl back}}\right) & \text{if } t > 0 \text{ and } x_{\text{nucl front}} \le x \le x_{\text{nucl back}} \\ 0 & \text{otherwise.} \end{cases} \tag{A.1}$$

We assume that the diffusion coefficient of active YAP varies between the cytoplasm and the nucleus as a hyperbolic tangent,

$$D_{Y_{\text{act}}} = D_{Y_{\text{nucl}}} + \left(D_{Y_{\text{cyto}}} - D_{Y_{\text{nucl}}}\right)\left[1 - \frac{1}{2}\left(\tanh\left[F(x - x_{\text{nucl front}})\right] - \tanh\left[F(x - x_{\text{nucl back}})\right]\right)\right], \tag{A.2}$$

where $F = 30$ is a scaling factor chosen to yield a transition region considerably narrower than the cell length.

Table A.1: Parameter definitions, values and units not listed in Table 1.

| Parameter | Meaning | Value / range | Units |
|---|---|---|---|
| $c_5$ | constant governing the feedback of YAP on Rho | $1 - 30$ | $\text{s}^{-1}C_{0\text{A}}$ |
| $c_6$ | base activation rate for YAP | 1 | $\text{s}^{-1}$ |
| $c_7$ | base deactivation rate for YAP | 1 | $\text{s}^{-1}$ |
| $c_8$ | Hill function parameter for YAP | $2.5 - 3.5$ | $C_{0\text{A}}^{-1}L$ |
| $c_9$ | Hill function parameter for YAP | 1 | 1 |
| $c_{10}$ | Hill function parameter for YAP | 5 | $C_{0\text{A}}^{-1}L$ |
| $q$ | effective charge of active YAP | $0.042 - 42$ | $\text{s}^{-1}C_{0\text{A}}^{-1}L^2$ |
| $q_0$ | effective charge of active YAP in the absence of a stimulus | $1.25 - 2.5$ | $\text{s}^{-1}L$ |
| $q_1$ | effective charge of inactive YAP | $0.5 - 2$ | $\text{s}^{-1}L$ |
| $Y_{\text{act nucl threshold}}$ | threshold value above which YAP deactivates Rho | $0.4 - 1$ | $N_{0\text{Y}}L^{-3}$ |
| $x_{\text{nucl front}}$ | $x$-coordinate of the front edge of the nucleus | 1/3 | $L$ |
| $x_{\text{nucl back}}$ | $x$-coordinate of the back edge of the nucleus | 2/3 | $L$ |
| $A_{\text{pert}}$ | amplitude of the initial parabolic perturbation | $10^{-2} - 1$ | 1 |
| $L_{\text{pert}}$ | width of the perturbed region (for a parabolic initial perturbation) | 0.15 | $L$ |
| $S_{\text{ampl}}$ | amplitude of the transient stimulus taken from Ref. (32) | $0.01 - 0.5$ | $\text{s}^{-1}$ |
| $t_1$ | duration of the application of a $t$-independent stimulus, see Ref. (32) | 20 | s |
| $t_2$ | duration of the application of a $t$-dependent stimulus, see Ref. (32) | 25 | s |

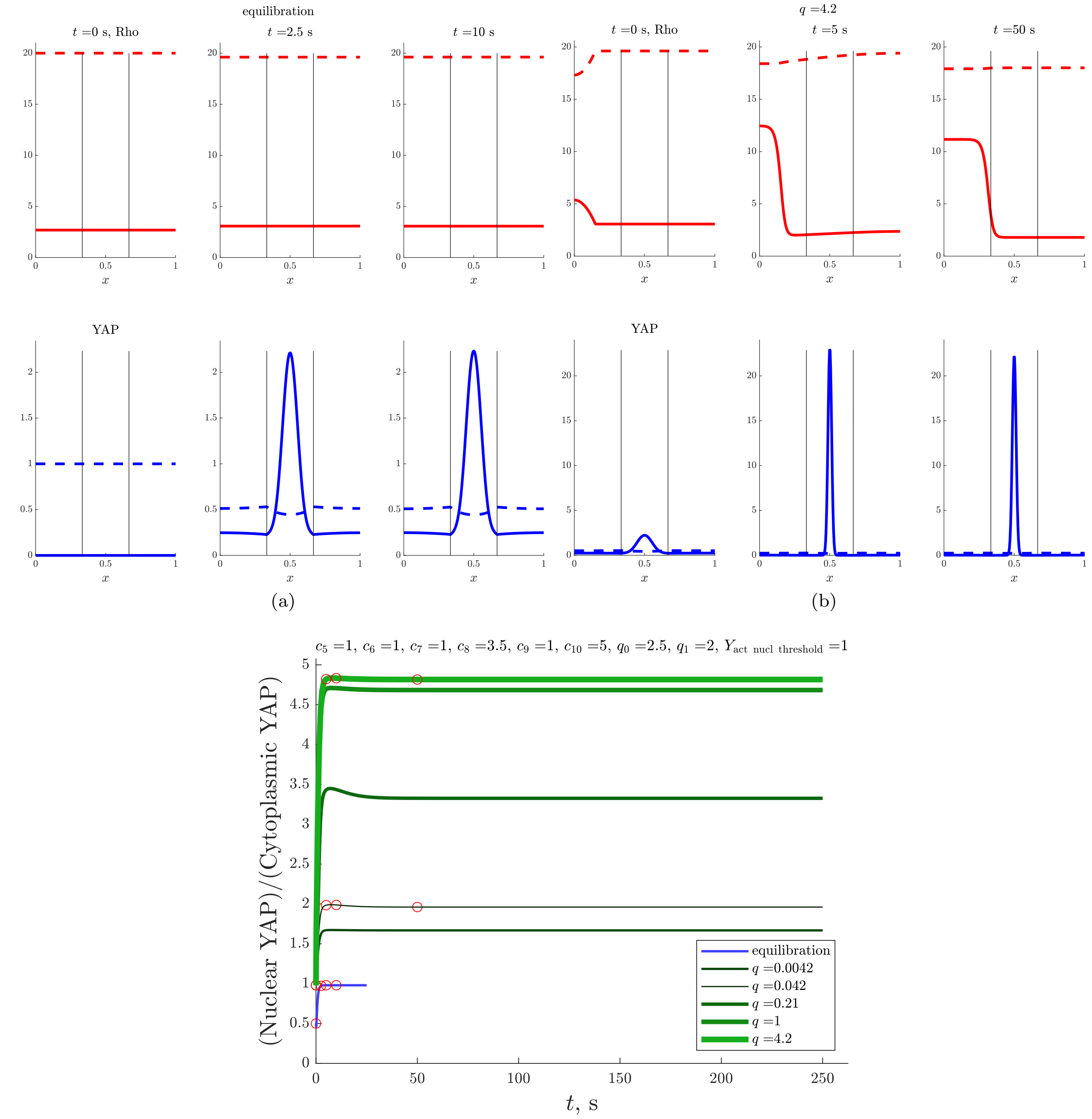


Figure A.1: **a**. Snapshots of the evolution of the concentrations of the different species in the system for an equilibration simulation with no perturbation in the Rho concentrations. Rho (top row) and YAP (bottom row), with solid lines depicting acting forms and dashed lines - inactive forms. The vertical lines indicate the boundaries of the nucleus. **b**. Snapshots of the evolution of the concentrations of the different species in the system at a high value of the mechanical forcing parameter $q$ - Rho (top row) and YAP (bottom row), with solid lines depicting acting forms and dashed lines - inactive forms. As seen in the plot for $t = 0\,\text{s}$, the initial perturbation to the homogeneous Rho concentrations is parabolic. The vertical lines indicate the boundaries of the nucleus. **c**. Evolution of the YAP ratio for simulations with a parabolic initial perturbation at different values of the mechanical forcing parameter $q$. Red circles mark the points in time at which the concentration profile snapshots are taken.

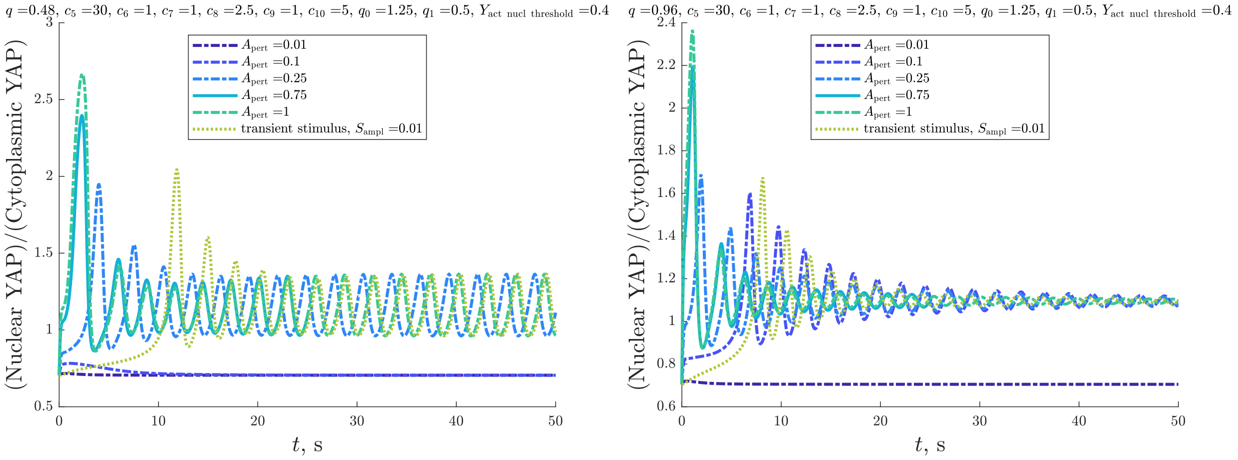


(a) (b)

Figure A.2: $R(t)$ for simulations with different initial conditions, namely parabolic perturbations of varying amplitudes (eq. (13)) and transient stimuli as per eqs. (14)-(15). The curves at $A_{\text{pert}} = 0.75$ are those plotted in Figure 3a; **a** illustrates the case of $q = 0.48$, which allows for sustained oscillations, whereas at $q = 0.96$, only damped oscillations are possible. For the simulations with a transient stimulus as per Ref. (32), we use a much lower $S_{\text{Ampl}}$ than the value of 0.5 in Figure 2a because the larger YAP-Rho coupling constant $c_5$ here would otherwise cause negative values of the concentrations of some species.

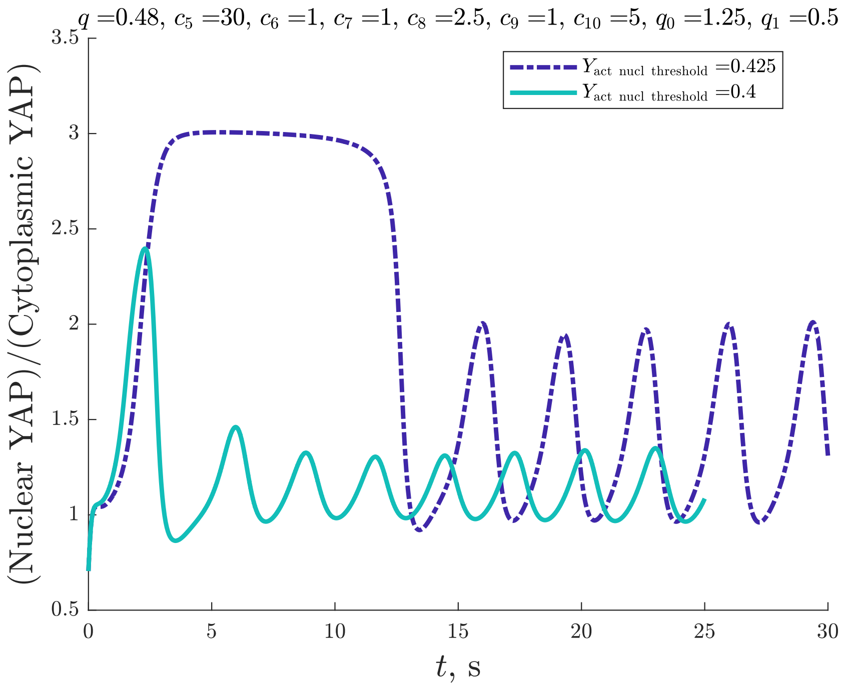


(a)

Figure A.3: $R(t)$ for a simulation with the same parameters as the one in Figure 3b except for $Y_{\text{act nucl threshold}} = 0.425$ rather than $Y_{\text{act nucl threshold}} = 0.4$. Note that $R(t)$ oscillates about $\approx 1.5$ rather than 1.15 as in Figure 3b.
